# Diacylglycerol metabolism drives host-pathogen responses during enteric infection in *Drosophila*

**DOI:** 10.1101/2025.08.28.672769

**Authors:** Xiaotong Li, Mohamed Mlih, Jason Karpac

## Abstract

Lipid metabolism is fundamental to cellular homeostasis, supporting energy storage, membrane architecture, and cellular signaling. Beyond these canonical roles, lipids have emerged as critical regulators of host immunity. Here, we define a lipid-driven mechanism that governs host-pathogen interactions by impacting pathogen clearance and thus infection outcomes. Exploiting Drosophila, we show that enteric infection triggers robust accumulation of neutral lipids, and specifically 1,2-diacylglycerols (DAGs), in the midgut. Disruption of DAG biosynthesis or lipid transport in midgut enterocytes (ECs) impairs lipid accumulation and reduces host survival. Conversely, dietary lipid supplementation enhances lipid storage and improves survival. Mechanistically, these lipid-dependent responses regulate defecation, thereby controlling bacterial clearance from the midgut. DAGs can act as signaling lipids that activate protein kinase C (PKC), and DAG accumulation in ECs correlates with elevated PKC activity and calcium signaling in midgut visceral muscle (VM), promoting VM contraction, midgut motility, and expulsion of pathogens via defecation. Together, our findings reveal a previously unrecognized role for DAG metabolism in shaping host defenses.

## Introduction

Lipid metabolism is central to cellular physiology, supporting diverse functions from energy homeostasis to membrane biogenesis to signal transduction^1–3^. Beyond these canonical roles, lipids are increasingly recognized as active regulators of immune function, particularly in the context of host-pathogen interactions^4^. Lipids can modulate immune responses by regulating signaling pathways, altering membrane properties, and serving as bioactive mediators that shape susceptibility to infection^5,6^. These functions are especially critical during pathogenic bacterial infections, where lipid metabolic remodeling may either support pathogen clearance or contribute to disease progression^7,8^. There is thus a critical need to explore the mechanistic basis through which distinct lipid metabolic cycles and lipid species govern host-pathogen dynamics and infection outcomes.

Dietary inputs represent a major determinant of this lipid-dependent immuno-metabolic balance, linking nutrient and metabolite availability to the capacity of the host to conduct effective defense against infectious agents^9^. For example, macronutrients such as glucose, amino acids, and lipids are sensed by systemic and tissue-resident immune cells, thereby shaping their differentiation, effector function, and survival^10–14^. Furthermore, nutrient surplus, present in obesity and instigated by Western-type diets, promotes chronic low-grade inflammation and a range of metabolic diseases which compromise host defense against pathogens^15,16^. Caloric restriction, conversely, can enhance resilience and modulate immune memory, in some cases improving pathogen tolerance^17,18^. Macronutrients like lipids thus play a crucial role in tuning immunity and promoting either physiological or pathophysiological responses to infection.

To this end, it has been increasingly recognized that specific lipid classes and species exert divergent effects on both innate and adaptive immune pathways^19^. Saturated fatty acids such as palmitate activate pro-inflammatory signaling cascades, including inflammasome pathways, whereas unsaturated fatty acids, particularly omega-3 fatty acids like DHA and EPA, dampen cytokine production and promote resolution of inflammation^20,21^. Cholesterol content modulates membrane raft composition and antigen receptor signaling, influencing both immune cell activation and innate immune responses^22^. Sphingolipids, including ceramides and sphingosine-1-phosphate, regulate immune cell trafficking, polarization, and survival, thereby linking lipid flux to inflammatory phenotypes^23^. Notably, diacylglycerols (DAGs) serve as second messengers that activate protein kinase C (PKC) and downstream transcriptional programs critical for lymphocyte activation and antimicrobial defense^24,25^. These findings underscore that distinct lipid species act as immuno-regulatory signals, and the balance of metabolic cycles that dictate the anabolism and catabolism of these lipids plays a critical role in defining immune responses. Furthermore, because distinct tissue-types have unique metabolic features, lipid-immune integration is likely to be defined by cellular specificity.

As a primary site of host–microbe interactions in most animals, the gastrointestinal tract represents a metabolically and immunologically dynamic environment^26,27^. The gastrointestinal tract of many invertebrates (including insects) also serves a primary barrier epithelium to external bacterial pathogens, making these tissues excellent models for exploring ancient immuno-metabolic defense mechanisms. In the fly *Drosophila melanogaster*, the midgut (a primary component of the gastrointestinal tract) serves a functionally analogous role to the vertebrate intestine with primary immune-, metabolic-, and nutrient (diet)-dependent functions^28^. Midgut epithelial enterocytes (ECs) absorb dietary lipids, synthesize neutral lipids for storage in lipid droplets, and distribute systemic lipids via lipophorins^29^. Conserved lipid metabolic regulators, such as sterol regulatory element-binding protein (SREBP), target of rapamycin (TOR), and insulin signaling integrate nutritional cues to shape lipid anabolism and catabolism^30–33^. Concurrently, the midgut epithelium serves as an immune barrier, detecting symbiotic and pathogenic microbes and activating local and systemic innate immune responses through conserved pathways controlling innate immune regulators such as nuclear factor kappa-light-chain enhancer of activate B-cells (NFkB), dual oxidase (Duox), and reactive oxygen species (ROS)^34,35^. Enteric pathogens disrupt midgut homeostasis and trigger metabolic rewiring that impacts host resilience^36,37,28^. Thus, the Drosophila midgut offers a powerful model to study the interplay between lipid metabolic cycles and innate immunity that direct host-pathogen responses in a genetically tractable system^28,38,39^.

In this study, we leverage the Drosophila midgut to dissect the specific roles of lipids and lipid metabolic cycles in modulating host infection outcomes during enteric infection. Specifically, we examine how distinct lipid species, their biosynthetic pathways, and lipid transport coordinate physiological responses that promote host survival. Our findings reveal a lipid-dependent signaling axis that links midgut epithelial cells and midgut muscle to midgut motility and bacterial pathogen clearance, offering mechanistic insight into how lipid dynamics orchestrate host defenses.

## Results

### Enteric infection leads to neutral lipid accumulation in the Drosophila midgut

To examine the tissue-autonomous impact of enteric infection on host lipid metabolism, we utilized a previously characterized Drosophila model in which adult flies are orally infected with *Pseudomonas entomophila* (*P.e.*) for 20 hours at 25 °C^40^. *P.e.* is a natural entomopathogen^41^. Fluorescent staining using Nile Red and BODIPY 493/503 revealed a marked accumulation of neutral lipids in the midgut of wild-type (OreR) flies during infection, consistent with increased lipid droplet (LD) content (Fig. 1A–D). Neutral lipids accumulate throughout the midgut in response to enteric infection, with a pronounced increase in the posterior midgut (Fig. 1A-D). Additionally, CholEsteryl BODIPY staining further demonstrated enhanced cholesterol levels during *P.e.* infection, suggesting broad remodeling of lipid metabolism within the midgut intestinal epithelium (Fig. S1A, B).

**Figure 1.**
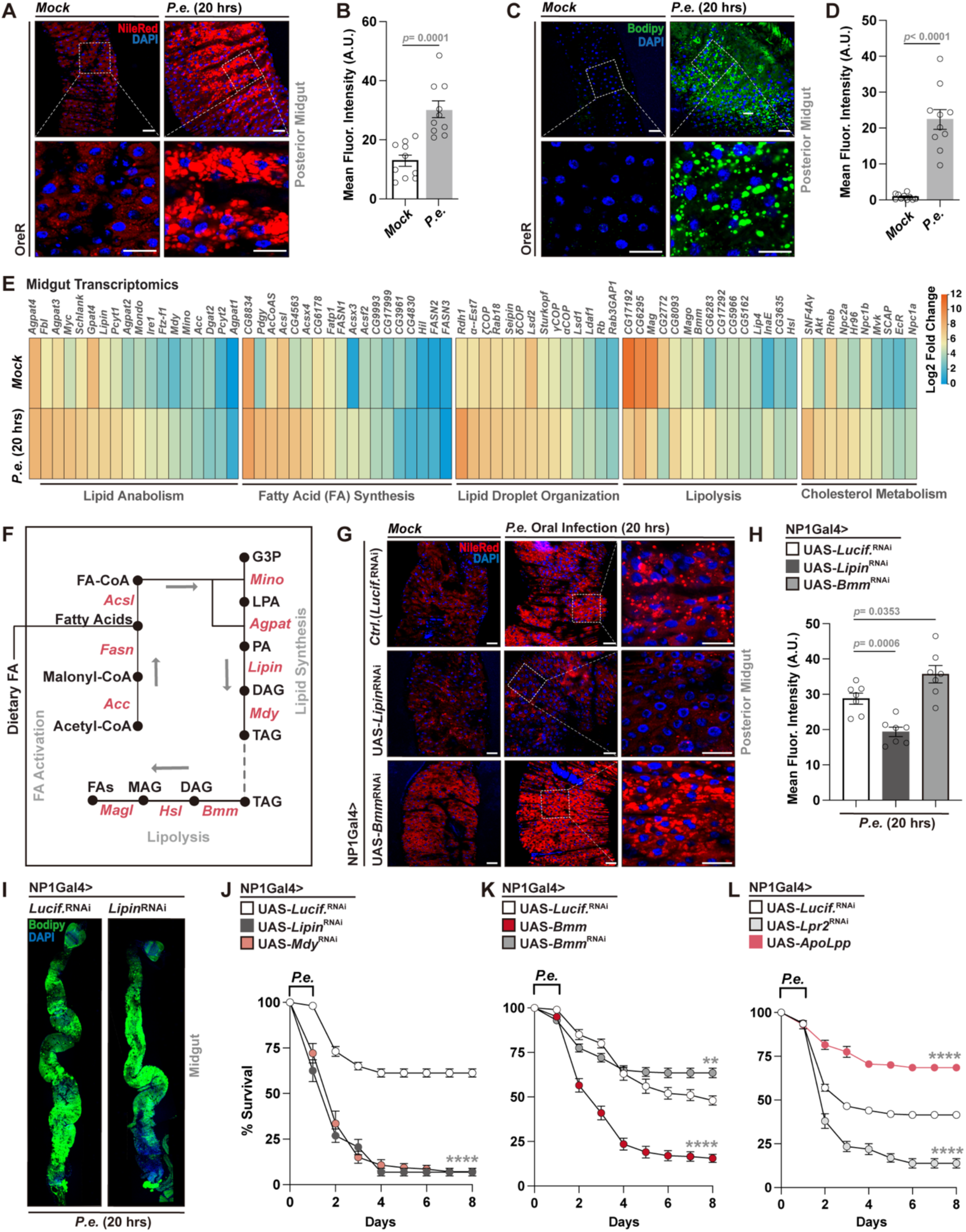
Enteric infection with *P.e.* induces lipid accumulation in the Drosophila midgut and modulates host survival. (A, B) Nile Red staining (A) and intensity quantification (B) of neutral lipids in the posterior midgut (genotype; OreR) during mock treatment or *P.e.* infection (20 hours after inoculation/initial feeding; n = 10 samples). Lipid droplet (Nile Red, red), and nuclei (DAPI, blue). (C, D) Bodipy493/503 staining (C) and intensity quantification (D) of neutral lipids in the posterior midgut (OreR) during mock treatment or *P.e.* infection (n = 10 samples). Lipid droplet (Bodipy 493/503, green), and nuclei (DAPI, blue). (E) RNA-seq heatmap showing lipid metabolic-related gene expression in midguts during *P.e.* infection (compared to mock treatment; genotype – NP1Gal4>UAS-*Lucif.*^RNAi^). (F) Simplified lipid metabolic pathway highlighting manipulated enzymes. (G, H) Nile Red staining (G) and intensity quantification (H) of neutral lipids in posterior midgut from flies with EC-specific (NP1Gal4) attenuation of Lipin or Bmm during *P.e.* infection (n = 7 samples). Lipid droplet (Nile Red, red), and nuclei (DAPI, blue). (I) Bodipy493/503 staining showing reduced lipid accumulation in NP1Gal4>UAS-*Lipin*^RNAi^ posterior midgut during infection. Lipid droplet (Bodipy 493/503, green), and nuclei (DAPI, blue). (J) Survival curves post-infection following EC-specific attenuation of lipid synthesis genes (NP1Gal4>UAS-*Lipin*^RNAi^ or UAS-*Mdy*^RNAi^) compared to controls (NP1Gal4>UAS-*Lucif.*^RNAi^). n = 200 flies per genotype. (K) Survival curves post-infection following genetic manipulation of Bmm in ECs (NP1Gal4>UAS-*Bmm* or UAS-*Bmm*^RNAi^) vs. control (NP1Gal4>UAS-*Lucif.*^RNAi^). n = 200 flies per genotype. (L) Survival curves post-infection following genetic manipulation of lipid transportation genes in ECs (NP1Gal4>UAS-*ApoLpp* or UAS-*Lpr2*^RNAi^) compared to controls (NP1Gal4>UAS-*Lucif.*^RNAi^). n = 200 flies per genotype. Statistical comparisons were performed using two-sided t-tests (B, D, H) or two-way ANOVA (J-L). Bars represent mean ± SEM. **p value < 0.01, ****p value < 0.0001. Scale bars, 10 μm.

In order to begin to define the molecular basis of these changes, we performed RNA sequencing of whole dissected midguts from mock- and *P.e.*-infected flies. Transcriptomic profiling revealed robust upregulation of genes involved in lipid anabolism, fatty acid biosynthesis, LD organization, and cholesterol metabolism, alongside suppression of genes associated with lipolysis (Fig. 1E). These findings were validated by qPCR, which confirmed increased expression of key lipogenic genes including *Fasn1*, *Lipin*, and *Mdy* (Fig. 1F, S1C). Together, these results indicate that enteric *P.e.* infection induces a coordinated transcriptional program that promotes lipid uptake, synthesis, and storage in the Drosophila midgut.

### Enterocyte lipid metabolism modulates host infection outcomes

To explore the lipid metabolic cycles and pathways underlying infection-induced lipid accumulation, we genetically perturbed lipid metabolism specifically in ECs, which orchestrate local and systemic lipid homeostasis via absorption, synthesis, and mobilization (Fig. 1F). EC-specific attenuation of *Lipin* (NP1Gal4>UAS-*Lipin*^RNAi^), encoding an enzyme that converts phosphatidic acid (PA) to diacylglycerol to promote lipid anabolism (Fig. 1F), markedly reduced lipid levels in the posterior midgut, as shown by Nile Red and BODIPY 493/503 staining (Fig. 1G–I). In contrast, attenuation of the triglyceride lipase *Brummer* (*Bmm,* a critical regulator of lipolysis and lipid catabolism) resulted in elevated lipid accumulation during *P.e.* infection (Fig. 1F–H). We also found additional genetic interventions to modulate midgut lipid storage during enteric infection through overexpressing *apolipophorin* (*ApoLpp*). ApoLpp is a critical inter-cellular transporter of neutral lipids in Drosophila^42,43^, and while the gene is not endogenously expressed in cells of the midgut^42^, exogenous overexpression of *ApoLpp* in midgut ECs (NP1Gal4>UAS-*ApoLpp*) stimulates lipid storage and boosts infection-mediated lipid accumulation (Fig. S1D). Finally, suppression of *Srebp*, a master regulator of lipid synthesis and uptake of lipids^44^, led to decreased neutral lipid levels in ECs during *P.e.* infection (Fig. S1E). In totality, these findings confirm that the regulation and balance of lipid anabolism, catabolism, and potentially transport in midgut ECs can tune intestinal lipid accumulation during enteric infection.

We next asked whether these lipid metabolic changes impact infection outcomes. Midgut EC-specific attenuation of lipid synthesis (anabolism) genes, *Lipin* or *Mdy*, (NP1Gal4>UAS-*Lipin*^RNAi^ or UAS-*Mdy*^RNAi^) significantly compromised survival of flies following *P.e.* challenge (Fig. 1J). Attenuation of *Bmm* (and limiting lipid catabolism) enhanced survival, while *Bmm* overexpression (and promoting lipid catabolism) had the opposite effect (Fig. 1K). Exogenous overexpression of *ApoLpp* (NP1Gal4>UAS-ApoLpp), which enhances lipid accumulation, also improved host survival outcomes. Conversely, disrupting the transport of lipids (from other tissues) into midgut ECs by attenuating *lipophorin receptor 2* (*Lpr2*) function (NP1-Gal4>UAS-*Lpr2*^RNAi^) impaired survival (Fig. 1L). Lpr2 is a transmembrane receptor expressed in many Drosophila tissues that is responsible for the cellular uptake of neutral lipids from apolipophorins^45^. Modulation of *Srebp* expression in ECs (and thus midgut lipogenesis) also bidirectionally altered survival outcomes following infection (Fig. S1F). Similar trends were observed using an independent EC genetic driver (MexGal4; MexGal4>UAS-*Lipin*^RNAi^, UAS-*Bmm*^RNAi^ or UAS-*Acc* [acetyl Co-A carboxylase, see Fig. 1F]), further validating these lipid metabolic-dependent effects on survival following infection (Fig. S1G, H).

Collectively, these results demonstrate that midgut EC-intrinsic lipid metabolism modulates both infection-mediated lipid accumulation and host survival. More specifically, metabolic shifts towards lipid anabolism and uptake in the midgut are required to promote host survival. These data thus provide a functional link between midgut lipid metabolism and host-pathogen responses that underly infection outcomes.

### Dietary lipid supplementation promotes midgut lipid accumulation during infection and enhances host survival

Having established that midgut EC-intrinsic lipid metabolism modulates infection outcomes, we next asked whether dietary lipid availability could similarly influence host-pathogen responses. To this end, Oleic Acid (OA), a monounsaturated fatty acid and lipid precursor for synthesis^46^, was incorporated at 5% into our standard Drosophila diet, and after five days of OA feeding flies were subjected to *P.e.* infection. OA supplementation significantly enhanced lipid accumulation in the posterior midgut, as shown by increased Nile Red staining, particularly in infected flies (Fig. 2A, B). Moreover, OA-fed Drosophila exhibited improved survival following enteric infection (Fig. 2C).

**Figure 2.**
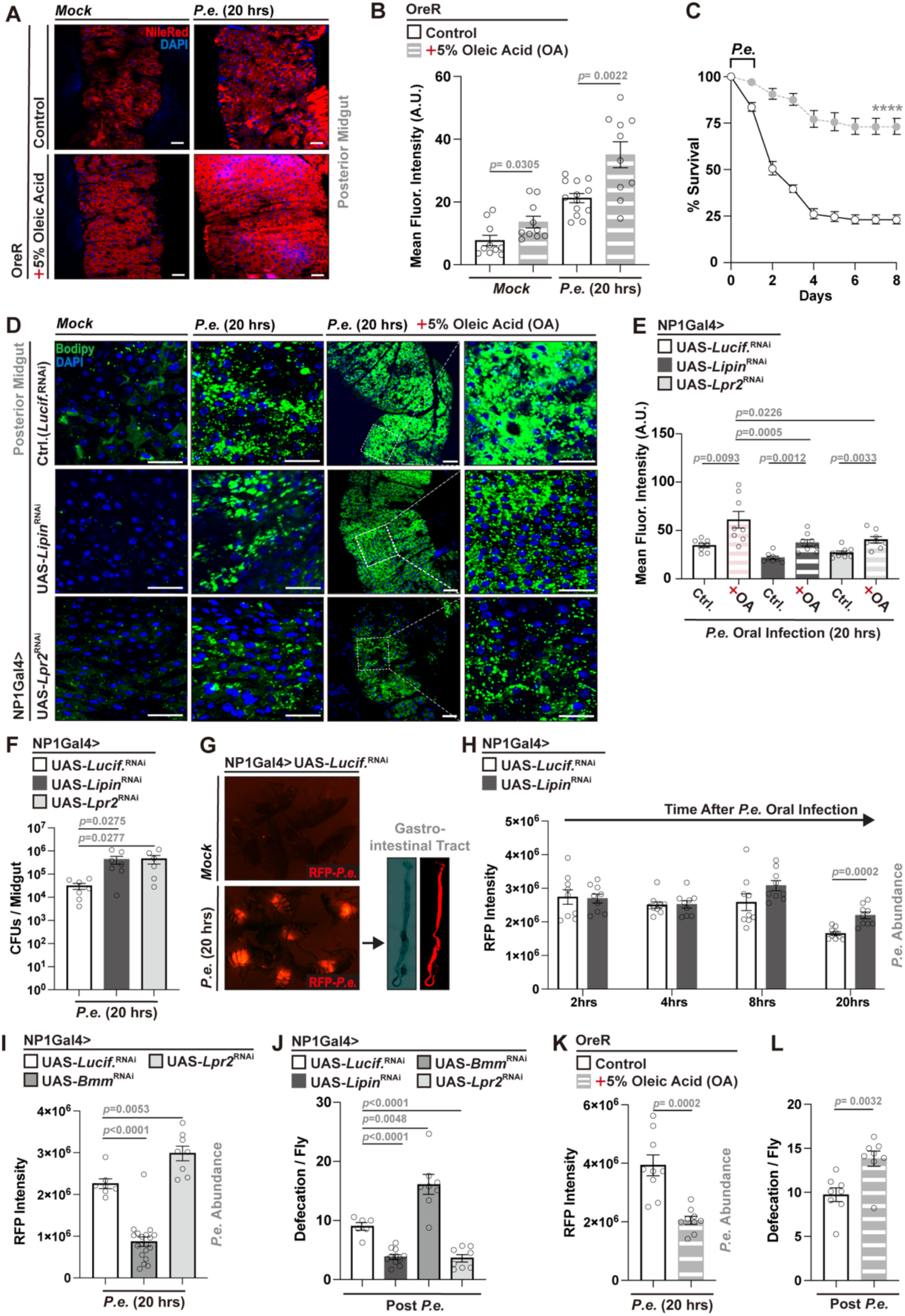
Dietary oleic acid (OA) supplementation and enterocyte lipid metabolism influence *P.e.*-dependent infection outcomes. (A, B) Nile Red staining (A) and intensity quantification (B) of posterior midgut lipid accumulation in flies (genotype; OreR) fed normal or 5% OA diets, with or without (mock) *P.e.* infection (n = 10). Lipid droplet (Nile Red, red), and nuclei (DAPI, blue). (C) Survival curves post *P.e.* infection comparing control vs. OA-fed flies. n = 200 flies per condition. (D, E) Bodipy493/503 staining (D) and intensity quantification (E) of lipid droplets in posterior midguts from NP1Gal4-driven control, *Lipin*, or *Lpr2* RNAi flies during mock treatment or during *P.e.* infection, ± OA diet (n = 8). Lipid droplet (Bodipy 493/503, green), and nuclei (DAPI, blue). (F) Gut bacterial load (CFUs) in flies with EC-specific (NP1Gal4) attenuation of *Lipin* or *Lpr2* during *P.e.* infection compared to controls (n = 7). (G) Representative images of whole flies and dissected intestines showing *P.e.-*RFP (red) accumulation after infection. (H, I) Quantification of RFP-expressing *P.e.* intensity in guts of NP1Gal4-driven *Lipin* (H) or *Bmm* and *Lpr2* (I) RNAi flies at indicated timepoints during infection (n = 7-20). (J) Defecation frequency per fly (16 h post-infection) in indicated RNAi genotypes (n = 6-11). (K, L) Effects of dietary OA supplementation on bacterial load (K, RFP intensity) and defecation frequency (L) (n = 8-9). Statistical comparisons were performed using two-sided *t*-tests (B, E, F, H-L) or two-way ANOVA (C). Bars represent mean ± SEM. ****p value < 0.0001. Scale bars, 10 µm.

Notably, OA supplementation could only partially restore lipid accumulation in midgut EC-specific *Lipin-* or *Lpr2*-attenuated flies during infection, highlighting that metabolic cycles and pathways required for lipid anabolism and transport are essential for infection-dependent lipid accumulation in the midgut, even when diets are supplemented with precursors for lipid synthesis (Fig. 2D, E). These findings indicate that dietary fatty acid availability modulates midgut lipid metabolism and impacts host survival during enteric pathogenic bacterial challenge.

### Midgut lipid metabolism regulates bacterial pathogen clearance through defecation

We next wanted to investigate the physiological mechanisms and outputs, driven by these adaptive midgut lipid metabolic responses, that could influence host infection outcomes. We first ruled out that deficits in host inflammatory responses (such as antimicrobial peptide [AMP] production) contributes to lipid-dependent changes in infection outcomes. Expression of canonical AMP genes (*Drs*, *Dro*, *CecA*, *Def*, *Atta*) was measured in dissected midguts of control and midgut EC-specific *Lipin*-attenuated flies, under both mock and *P.e.* infection conditions. Enteric infection induced robust AMP genes expression in both genotypes during *P.e.* infection (Fig. S2A), suggesting that impaired pathogen defense responses in *Lipin-*attenuated flies are not attributable to defective AMP induction.

Our prior work has shown that systemic (muscle-and adipose-driven) lipid mobilization is corelated with enhanced bacterial pathogen clearance (resistance) and elevated defecation^40^. We therefore hypothesized that lipid accumulation within the midgut epithelium promotes host defenses by stimulating gut motility, defecation, and expulsion of pathogenic bacteria to boost infection resistance. To rule out feeding behavior as a confounding factor, we first assessed food intake in flies with EC-specific attenuation of *Lipin* or *Bmm* and found no significant differences compared to controls (Fig. S2B). We then quantified bacterial load in dissected midguts following *P.e.* infection under conditions of genetically altered midgut EC lipid metabolism. Attenuation of *Lipin* or *Lpr2* in midgut ECs significantly increased midgut bacterial burden (correlating with decreases in host survival; Fig. 1J, L), as measured by colony-forming units (CFUs; Fig. 2F). To visualize pathogen dynamics *in vivo*, we generated a DsRed-expressing *P.e.* strain (RFP-*P.e.*) by introducing a pSW002-Pc-DsRed-Express2 plasmid. This *P.e.* strain exhibited comparable pathogenicity to the original *P.e.* strain, with distinct midgut fluorescence during infection, better enabling real-time tracking of pathogenic bacterial clearance (Fig. 2G, Fig. S2C). Using RFP intensity as a proxy for bacterial abundance, we tracked pathogen clearance during infection. Impaired bacterial clearance, following 20 hours infection, was evident in flies with EC-specific attenuation of *Lipin* (NP1Gal4>UAS-*Lipin*^RNAi^ or MexGal4>UAS-*Lipin*^RNAi^) (Fig. 2H, S2D). Similarly, silencing of *Bmm* or *Lpr2* in midgut ECs (NP1Gal4/MexGal4>UAS-*Bmm*^RNAi^, NP1Gal4>UAS-*Lpr2*^RNAi^) led to changes in RFP intensity, consistent with altered bacterial load, highlighting that enhancing lipid accumulation (limiting catabolism) increases bacterial clearance while blocking lipid accumulation (limiting transport and uptake) decreases clearance (Fig. 2I, S2D). Attenuation of *Mdy* or *Srebp* in ECs produced similar effects (Fig. S2E), indicating that infection-induced midgut lipid accumulation inversely correlates with pathogen burden and positively correlates with host survival.

Given the established link between gut motility and bacterial expulsion during infection, we next examined whether midgut lipid metabolism impacts defecation. Flies with midgut EC-specific attenuation of *Lipin* or *Lpr2* exhibited reduced defecation rates during infection, whereas attenuation of *Bmm* led to an increased frequency of defecation in response to infection (Fig. 2J). These results were validated using the MexGal4 genetic driver and targeting *Lipin*, *Bmm* or *Acc* in midgut ECs (Fig. S2F). Additionally, dietary OA supplementation decreased midgut bacterial load and enhanced defecation during infection compared to flies fed a standard (control) diet (Fig. 2K, L), phenocopying genetic-dependent lipid accumulation.

Together, these findings indicate that midgut EC lipid metabolism modulates gut motility and pathogenic bacterial clearance through maintaining defecation during enteric infection, providing a mechanistic link between infection-induced lipid accumulation and host defense responses.

### 1,2-Diacylglycerols (DAGs) accumulate during enteric infection and correlate with changes in midgut visceral muscle function

Thus far, our data show that adaptive midgut lipid metabolic responses play an important role in shaping host defenses against enteric bacterial infection. Specifically, enhanced lipid anabolism and uptake/transport in the midgut promotes bacterial pathogen clearance through maintaining gut motility and defecation to improve host survival. We next aimed to explore distinct mechanisms by which midgut lipid metabolism influences defecation through identification of specific lipid classes or species that accumulate in the midgut following enteric infection. To this end, we performed unbiased lipidomic profiling on whole dissected midguts from mock and *P.e.*-infected flies. Across two control genetic backgrounds (OreR and NP1Gal4>UAS-*Lucif.*^RNAi^), we found that *P.e.* infection led to elevated levels of 1,2-diacylglycerols (1,2-DAGs; Fig. 3A-B and Fig. S3A). Other lipid species or classes, such as triglycerides (TAGs) and acylcarnitines did not show a comparable level of induction, independent of their abundance in the midgut (example in Fig. 3A). Additional lipid species were only detected at very low levels in whole midgut samples (such as ceramides and sphingolipids), and infection-mediated induction of 1,3-diacylglycerols (1,3-DAGs) was less abundant and more variable than the induction of 1,2-DAGs (Fig. 3A and Fig. S3A-B). In totality, our data show that neutral lipids and lipid droplets accumulate in the midgut epithelium during enteric infection, with a particular enrichment in 1,2-diacylglycerol lipid species.

**Figure 3.**
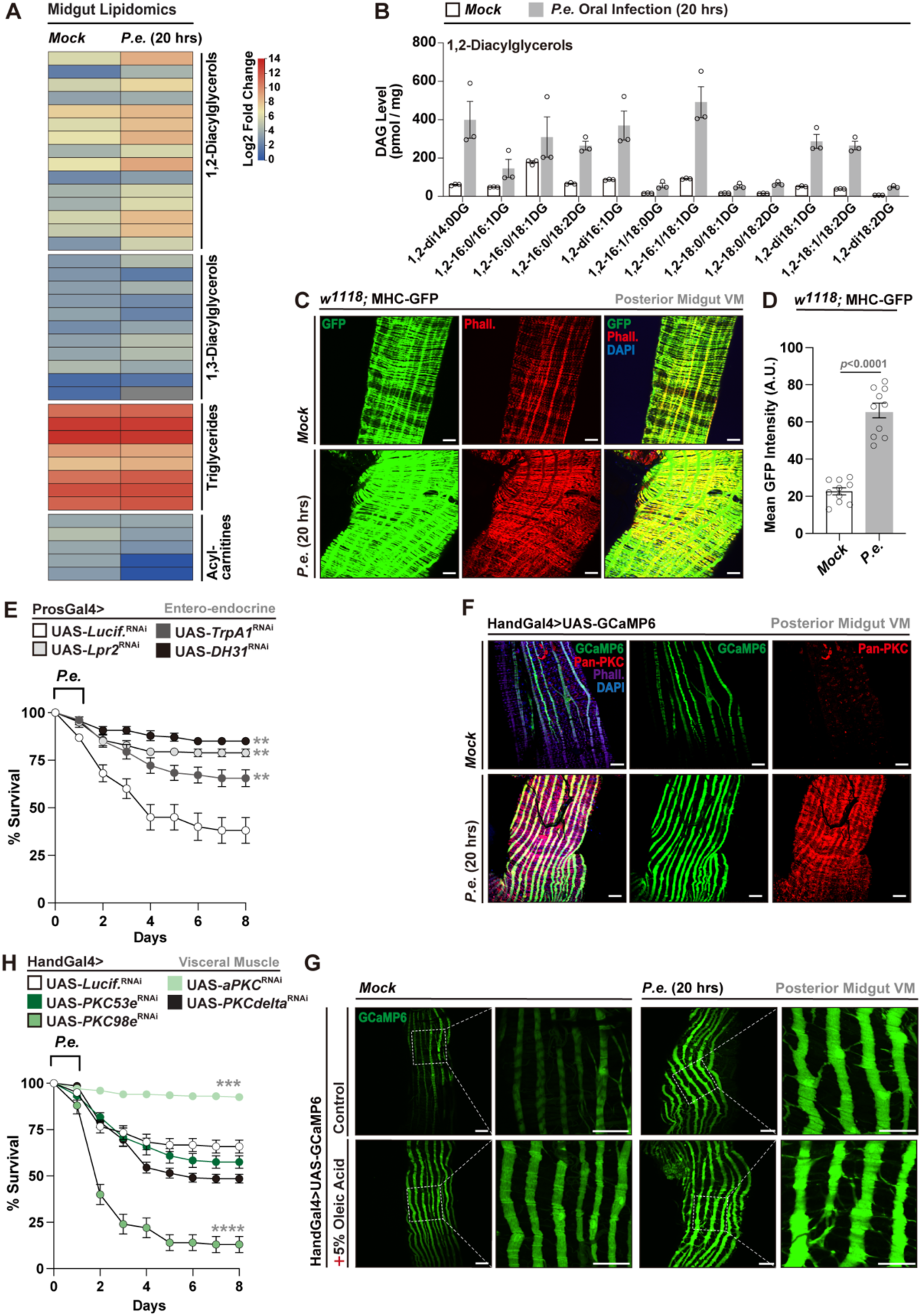
Enteric *P.e.* infection induces 1,2-DAG accumulation and midgut visceral muscle activation. (A, B) Lipidomic heatmap (A) and quantitative analysis (B) showing elevated 1,2-DAGs in whole fly midguts during *P.e.* infection (compared to mock controls). Heatmap representation of lipid species in dissected midguts from control (Mock) and *P. e.*–infected flies (20 h post-infection). Genotype in (A) – OreR; Genotype in (B) -NP1Gal4>UAS-*Lucif.*^RNAi^. Distinct lipid classes are indicated, including 1,2-diacylglycerols, 1,3-diacylglycerols, triglycerides, and acylcarnitines. (C, D) Increased visceral muscle activity during infection visualized by MHC-GFP fluorescence (green) and phalloidin staining (red, C), MHC-GFP intensity was quantified in (D) (n = 10). (E) Survival curves of flies with enteroendocrine cell-specific attenuation (RNAi) of *TrpA1*, *Lpr2* or *DH31* coding genes (ProsGal4> UAS-*TrpA1*^RNAi^, *Lpr2* ^RNAi^ or *DH31* ^RNAi^) post *P.e.* infection. n = 200 flies per genotype. (F, G) Calcium signaling activation in visceral muscle (HandGal4>UAS-GCaMP6; green) during infection (F), enhanced by OA supplementation (G). (H) Survival curves post-infection with visceral muscle (HandGal4) attenuation of *PKC* isoforms (target by RNAi). n = 200 flies per genotype. Statistical comparisons were performed using two-sided *t*-tests (B, D) or two-way ANOVA (E, H). Bars represent mean ± SEM. **p value < 0.01, ***p value < 0.001, ****p value < 0.0001. Scale bars: 10 μm.

Overall, the cellular pool of 1,2-DAGs can be generated through bulk lipid anabolism/synthesis, intake (transport or diet), and lipolysis of stored triglycerides (all of this is described in Figures 1 and 2), as well as more distinct mechanisms by which membrane phospholipids are cleaved into inositol trisphosphate (IP3) and 1,2-DAGs through phospholipase C (PLC) enzymes. Drosophila NorpA is a critical PLC-beta enzyme that generates 1,2-DAGs from phospholipids, while InaE (a DAG lipase) acts downstream of NorpA to breakdown DAGs^47,48^. We found that attenuating DAG synthesis or breakdown in midgut ECs via these mechanisms (NP1Gal4> UAS-*NorpA*^RNAi^ or UAS-*InaE*^RNAi^, respectively) also influenced host survival following *P.e.* infection. Attenuating *NorpA* in midgut ECs (limiting 1,2-DAG synthesis) decreased survival while attenuating *InaE* (blocking 1,2-DAG breakdown) enhanced survival (Fig. S3C). Our data, coupled with previous work, suggest that the Drosophila midgut may act as a major source of medium-chain DAGs^42,43^, and that midgut DAG metabolism might play a critical role in host-pathogen responses to enteric infection.

### Midgut enterocyte-derived DAG accumulation correlates with elevated PKC– calcium signaling activity in midgut visceral muscle during enteric infection

Unlike neutral lipids that are traditionally utilized for storage (such as 1,3-DAGs or TAGs), 1,2-DAGs function as potent signaling molecules. These DAGs can activate specific isoforms of protein kinase C (PKCs) through evolutionary conserved DAG sensing protein domains, and PKCs act as molecular phosphorylation switches to control various cellular functions across tissue-types^49,50^. Additionally, DAG metabolism, both indirectly and directly, can influence cellular calcium stores and calcium-dependent signaling pathways (including PKCs)^51^. The signaling integration of 1,2-DAGs, PKC, and calcium is known to control muscle function in a variety of muscle types, including muscle contraction and remodeling. Thus, we next asked whether DAG/lipid accumulation in midgut enterocytes contributes to host-pathogen responses by regulating gastrointestinal motility (and defecation/pathogen expulsion) via the midgut visceral muscle. The Drosophila midgut visceral muscle, like in most insects, is a contractile network surrounding the epithelium that is essential for peristalsis. This muscle is made up of both circular and longitudinal segments. Our whole midgut RNA-seq data revealed that enteric *P.e.* infection promotes the upregulation genes associated with muscle contraction and remodeling, including multiple *Actin* and *Myosin* isoforms, and other key regulatory components (Fig. S3D). Notably, expression of *Myosin heavy chain* (*MHC*), a key component of the contractile apparatus, was strongly induced. Utilizing *MHC*-GFP transcriptional reporter flies, we found that *P.e.* infection led to increased MHC-GFP intensity, as well as enhanced phalloidin (F-actin) staining, in midgut visceral muscle, indicating infection-induced activation and/or remodeling of the visceral musculature (Fig. 3C, D). These findings highlight that contractile remodeling of the midgut musculature is corelated with lipid/DAG accumulation during enteric infection.

Based on previous work in Drosophila, we hypothesized that visceral muscle contractility during enteric infection could be regulated by two distinct DAG-dependent mechanisms: First, a neuroendocrine route in which DAGs stimulate neuropeptide release from midgut enteroendocrine secretory cells (EECs) via activation of the TrpA1 receptor, or second, a paracrine route where DAGs produced in midgut ECs are transported to visceral muscle to activate a PKC–calcium signaling cascade^52–55^. To explore potential pathways that mediate host-pathogen responses during *P.e.* infection, we performed small, cell/tissue-specific RNAi screens targeting key components of each axis.

Disruption of the TrpA1–neuropeptide (DH31, a Drosophila secreted factor) pathway in EECs (ProsGal4>UAS-*TrpA1*^RNAi^ or UAS-*DH31*^RNAi^) did not decrease survival or bacterial clearance during *P.e.* infection (Fig. 3E, Fig. S3E). Similarly, attenuation of *Lpr2* (lipid transport) in EECs had no detrimental impact on infection outcomes (Fig. 3E). These findings suggest that a DAG–TrpA1–DH31 signaling route is dispensable for pathogen clearance in the context of enteric *P.e.* infection.

In contrast, we observed robust activation of visceral muscle calcium flux in *P.e.*-infected flies expressing GCaMP6, a widely used calcium indicator^56^, under a muscle-specific driver (HandGal4, enriched in visceral muscle), indicating enhanced contractile activity (Fig. 3F). This was accompanied by elevated pan-PKC protein levels, consistent with putative DAG-induced PKC signaling activity (Fig. 3F). Dietary oleic acid further increased GCaMP6 signal intensity under both baseline and infection conditions, reinforcing the link between midgut lipid abundance and visceral muscle activation (Fig. 3G).

To identify specific PKC isoforms mediating this response, we screened RNAi lines targeting multiple typical (*PKC53e*, *PKC98e*, *PKCδ*) and atypical (*aPKC,* lacking one of the conserved DAG-sensing domains) PKC family members (Fig. 3H). Among these, *PKC98e* attenuation resulted in the most pronounced reduction in survival following enteric infection (Fig. 3H, Fig. S4A), implicating it as a potential key effector of infection-mediated midgut visceral muscle remodeling/activation.

### Midgut visceral muscle PKC–calcium signaling and lipid transport are essential for pathogen clearance and shape infection outcomes

Together, our data suggest that midgut EC lipid/DAG accumulation shape midgut visceral muscle function, through PKC98e-calcium signaling, to enhance gut motility and facilitate pathogen expulsion. To mechanistically validate the role of this signaling axis in visceral muscle, we genetically disrupted these pathways using tissue-specific RNAi. Muscle-specific knockdown of *PKC98e* or *Itpr* (inositol 1,4,5-trisphosphate receptor), a key mediator of calcium release from the endoplasmic reticulum^57^, impaired visceral muscle architecture and remodeling during *P.e.* infection (visualized through MHC immunostaining; Fig. 4A). These perturbations, performed using two different muscle genetic drivers (HandGal4 and HowGal4), resulted in increased bacterial loads, defective pathogen expulsion, and decreased defecation (Fig. 4B, C; Fig. S4B-E). Attenuation of *Itpr* also significantly reduced host survival in response to infection (Fig. 4D, Fig. S4F), similar to attenuation of *PKC98e*, highlighting a functional requirement for PKC–calcium signaling in host-pathogen responses to *P.e.* infection. Notably, dietary OA supplementation could not significantly rescue defects in defecation or host survival when *PKC98e* is attenuated in muscle (Fig. 4E, F). These data, coupled with findings provided in Fig. 2D-E, highlight that PKC function in visceral muscle is crucial for the ability of midgut EC lipid/DAG accumulation to shape midgut motility and infection outcomes.

**Figure 4.**
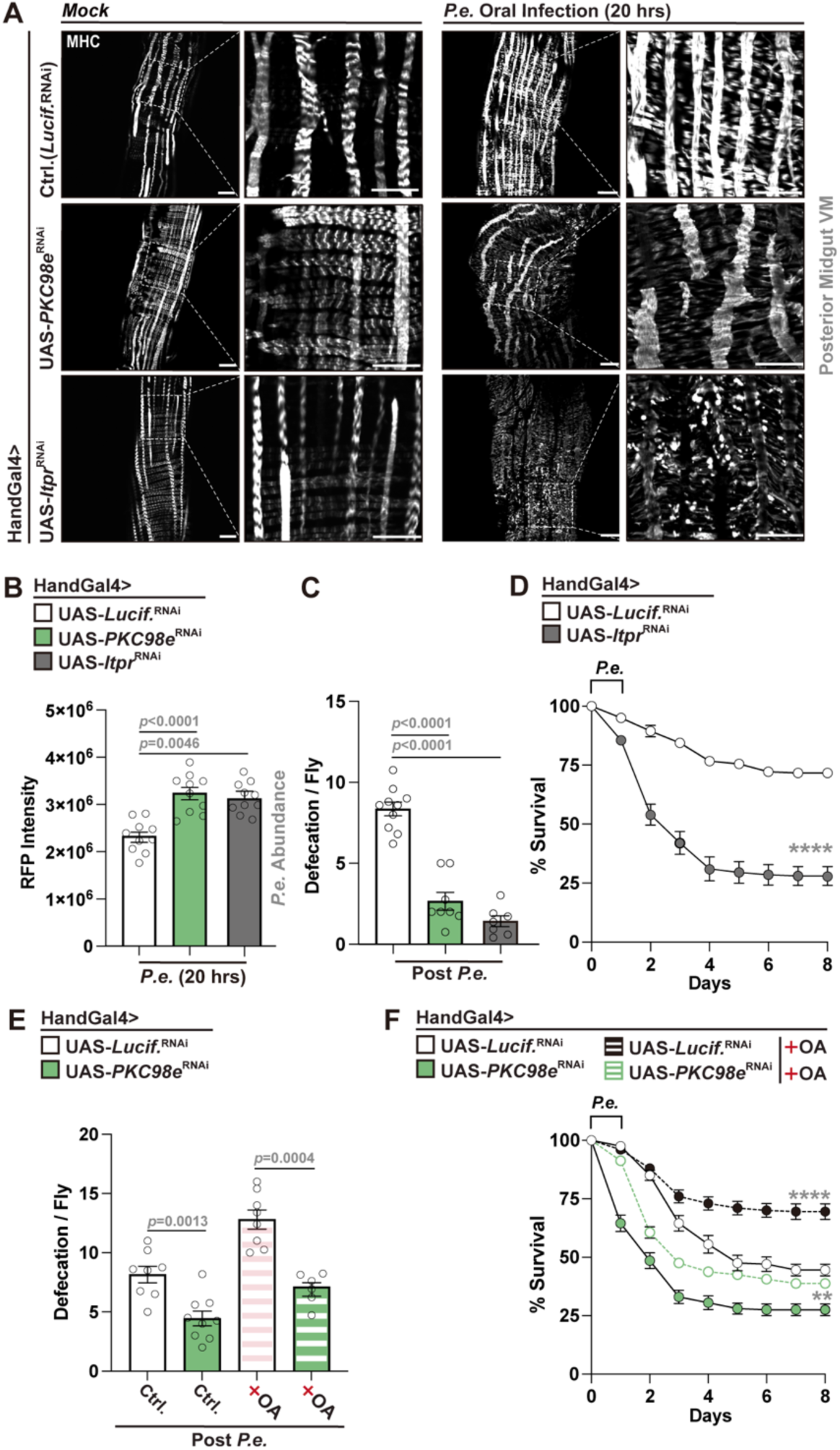
**PKC–calcium signaling in visceral muscle controls muscle remodeling, pathogen clearance, and host survival during *P.e.* infection**. (A) MHC immunostaining (white); indicaing muscle remodeling defects during *P.e.* infection using visceral muscle attenuation (HandGal4) of *PKC98e or Itpr* using RNAi. (B, C) Increased bacterial load (B, *P.e.-*RFP intensity) and decreased defecation (C) in HandGal4>UAS-*PKC98e*^RNAi^ or *Itpr*^RNAi^ flies compared to controls (NP1Gal4>UAS-*Lucif.*^RNAi^) post *P.e.* infection. (D) Survival curves for HandGal4>UAS-*Itpr*^RNAi^ flies compared to controls. n = 200 per genotype. (E, F) OA supplementation does not infleunce defecation (E) and survival (F) defects in HandGal4>UAS-*PKC98e*^RNAi^ flies. n = 8 in (E), n = 200 in (F). Statistical comparisons were performed using two-sided *t*-tests (B, C, E) or two-way ANOVA (D, F). Bars represent mean ± SEM. **p value < 0.01, ****p value < 0.0001. Scale bars: 10 μm.

We next explored the role of lipid transport in midgut visceral muscle during enteric infection. Attenuation of *Lpr2* in muscle (HandGal4>UAS-*Lpr2*^RNAi^) impaired visceral muscle architecture and remodeling during *P.e.* infection (Fig. 5A), and similarly disrupted pathogen clearance, defecation, and host survival (Fig. 5B-D). We also exploited the same genetic strategy that was used to enhance lipid transport in midgut ECs to promote lipid transport in midgut visceral muscle, specifically, exogenous activation of the lipophorin ApoLpp. Boosting lipid transport by overexpressing *ApoLpp* in visceral muscle significantly enhanced survival following enteric infection (Fig. 5E), altogether suggesting that lipid transport in midgut visceral muscle is required for muscle remodeling/contractility during infection. To this end, exogenous overexpression of *ApoLpp* in visceral muscle improved bacterial pathogen clearance and defecation following *P.e.* infection, but these benefits (including survival) were completely abolished by simultaneous attenuation of *PKC98e* (HowGal4>UAS-*ApoLpp*, UAS-*PKC98e*^RNAi^) (Fig. 5F–I), suggesting that lipid transport in visceral muscle requires PKC signaling activity to influence host-pathogen responses. These data collectively support a model in which enteric infection induces lipid/1,2-DAG accumulation in midgut ECs, and that midgut visceral muscle utilizes these lipids (through PKC-calcium signaling) to maintain gastrointestinal motility, promote pathogen expulsion, and ultimately impact infection outcomes.

**Figure 5.**
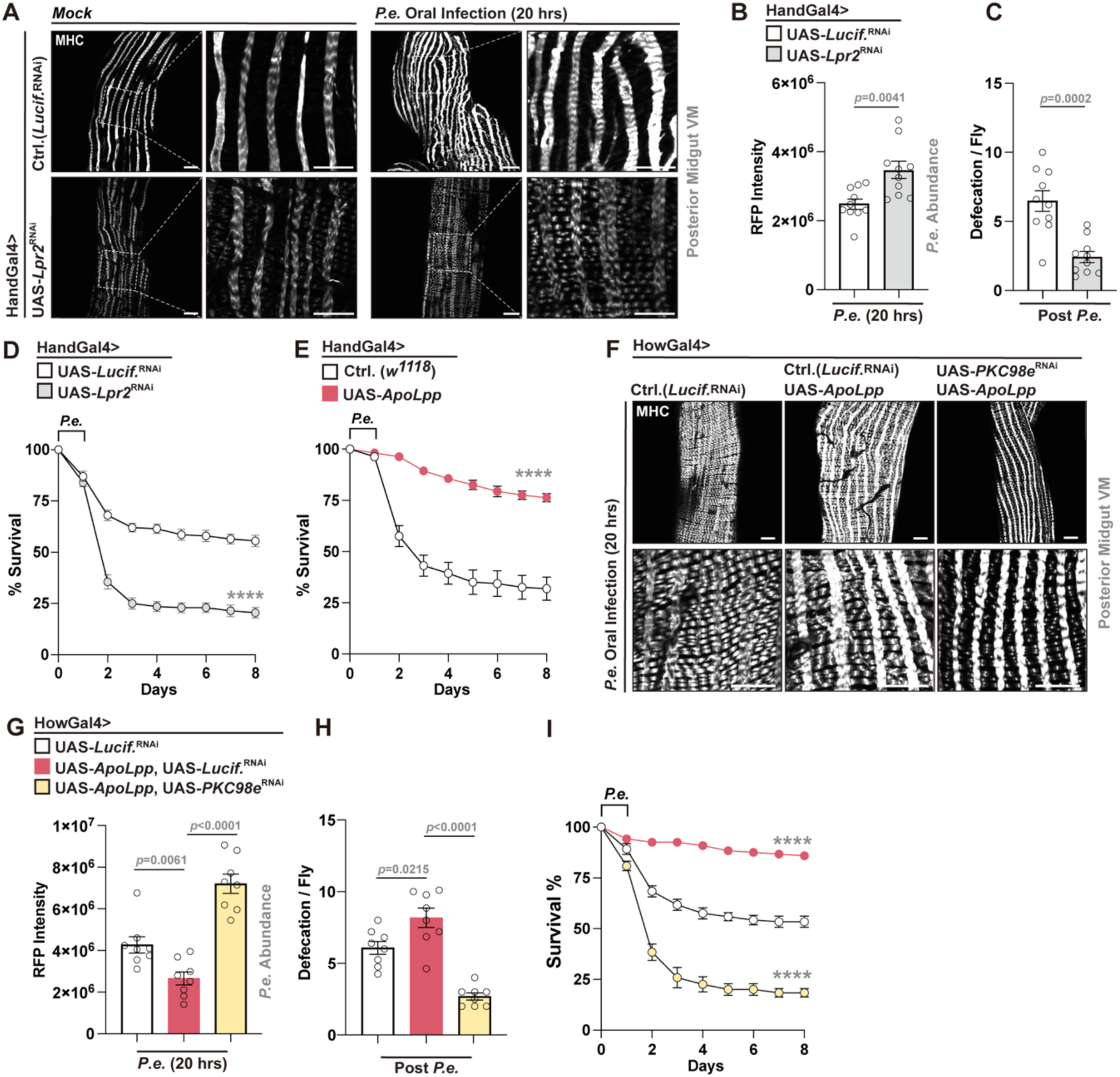
Lipid metabolism in visceral muscle controls muscle remodeling, pathogen clearance, and host survival during *P.e.* infection. (A) MHC immunostaining (white); indicating muscle remodeling defects during *P.e.* infection using visceral muscle attenuation (HandGal4) of *Lpr2* (RNAi). (B, C) Increased bacterial load (B, *P.e*-RFP intensity) and decreased defecation (C) in HandGal4>UAS-*Lpr2*^RNAi^ flies compared to controls (NP1Gal4>UAS-*Lucif.*^RNAi^) post *P.e.* infection. (D) Survival curves for HandGal4>UAS-*Lpr2*^RNAi^ flies vs. control. n = 200 per genotype. (E) Survival curves for HandGal4>UAS-*ApoLpp* flies compared to controls (HandGal4>*w1118*). n = 200 per genotype. (F-I) *ApoLpp* overexpression in visceral muscle (HowGal4) improves muscle contraction (F), pathogen clearance (G), defecation (H), and survival (I), and the effects are reversed by *PKC98e* attenuation (RNAi). n = 8 in (G, H), and n=200 per genotype in (I). Statistical comparisons were performed using two-sided *t*-tests (B, C, G, H) or two-way ANOVA (D, E, I). Bars represent mean ± SEM. ****p value < 0.0001. Scale bars, 10 µm.

### Tropomyosin1 regulates midgut visceral muscle contraction to control bacterial pathogen clearance and host survival

We next wanted to show, directly, that midgut visceral muscle contraction can shape defecation and regulate host-pathogen responses. Our midgut transcriptomics revealed that, among a multitude of muscle structure and/or function, Tropomyosin1 (Tm1) is induced during *P.e.* infection (Fig. S3D). Tm1 is a key component of actin filaments, a conserved regulator of muscle contractility, and has been implicated in PKC-and calcium-mediated muscle contraction^58,59^, (Fig. S3D). Visceral muscle-specific attenuation of *Tm1* (HandGal4>UAS-*Tm1*^RNAi^) disrupted muscle architecture and remodeling during enteric infection, suggesting defective muscle contractility (Fig. 6A). Functionally, *Tm1* attenuation using two independent genetic drivers (HandGal4 or HowGal4), led to increased midgut bacterial burden (reduced pathogen expulsion), decreased rates of defecation, and decreased host survival following *P.e.* infection (Fig. 6B–D, Fig. S5A–C). Moreover, improved pathogen clearance, defecation and host survival induced by exogenous *ApoLpp* overexpression in muscle were abolished by simultaneous attenuation of *Tm1* (HowGal4>UAS-*ApoLpp*, UAS-*Tm1*^RNAi^), leading poor host infection outcomes (Fig. 6E–G). Dietary OA supplementation prior to infection also could not rescue the defecation defects caused by *Tm1* knockdown during *P. e.* infection (Fig. S5D), indicating that Tm1 is a key effector linking midgut lipid metabolism to visceral muscle contractile function, gastrointestinal motility, and host-pathogen responses.

**Figure 6.**
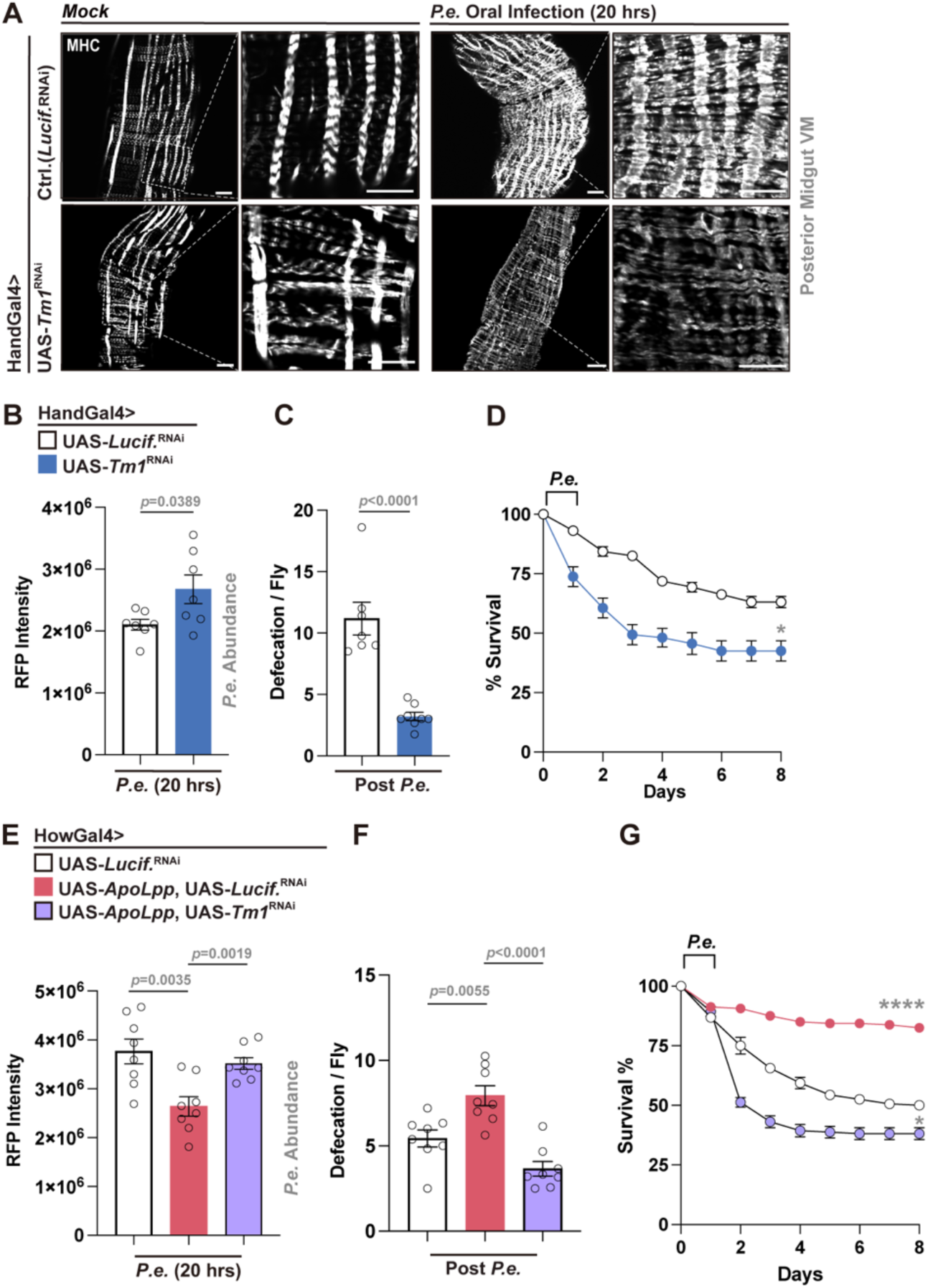
Tropomyosin1 (Tm1) is required to regulate infection-mediated visceral muscle contraction, pathogen clearance, and survival. (A) Visceral muscle (MHC immunostaining; white) remodeling defects in visceral muscle Tm1-attenuated flies (HandGal4>UAS-*Tm1*^RNAi^) during mock treatment or *P.e.* infection. (B, C) Increased bacterial load (B) and decreased defecation (C) in HandGal4>UAS-*Tm1*^RNAi^ flies compared to controls (n = 7-8). (D) Survival curves of visceral muscle-specific *Tm1*-attenuation flies compared to controls post *P.e.* infection. n=200 per genotype. (E–G) ApoLpp overexpression in visceral muscle (HowGal4>UAS-*ApoLpp*) improves bacterial clearance (E), defecation (F), and survival (G), and the effects are reversed by *Tm1* attenuation (HowGal4>UAS-*ApoLpp*, UAS-*Tm1*^RNAi^). n = 7-8 in (E, F). n=200 per genotype in (G). Statistical comparisons were performed using two-sided *t*-tests (B, C, E, F) or two-way ANOVA (D, G). Bars represent mean ± SEM. *p value < 0.05, ****p value < 0.0001. Scale bars, 10 µm.

## Discussion

Our study reveals a previously unrecognized role for diacylglycerol metabolism in shaping host susceptibility to enteric bacterial infection in Drosophila. By integrating lipidomic, transcriptomic, and functional genetics, we find that infection-induced lipid/1,2-DAG accumulation in the midgut can promote the activation of PKC–calcium signaling in visceral muscle to promote muscle contraction through Tropomyosin1 (Tm1). These signaling modalities enhance midgut visceral muscle remodeling, motility, and pathogen expulsion via increased defecation, thereby improving host survival. These findings also establish that distinct lipid metabolic cycles within the intestine can act as critical determinants of divergent host immune responses and further extend the functional repertoire of DAGs.

Lipid metabolism is increasingly recognized as a crux in shaping host-pathogen dynamics. Bacterial pathogens exploit host lipids as nutrients to support their growth and pathogenicity^60–62^. Conversely, the host mobilizes lipid-based defenses, including antimicrobial lipids, nutrient restriction, and immune cell modulation and activation^63–65^. Our findings add to this paradigm by demonstrating that tissue-specific and infection-induced lipid anabolism actively facilitates bacterial clearance, likely functioning in parallel with more canonical local and systemic immune responses, such as inflammation, AMP production, and/or generation of reactive oxygen species.

Drosophila models have provided fundamental insight into this interaction between lipid metabolism and immune responses. Previous studies revealed that septic bacterial infection of adult flies with *E. coli*, *P. asymbiotica* or *P. luminescens* resulted in dramatic accumulation of lipid droplets in the midgut^66^. Lipid accumulation is mediated through Gram-negative cell wall recognition machinery, and lipid accumulation can either provide resistance or be deleterious for the infected flies depending on the type of bacterial infection. However, another study showed gram-positive bacteria *E. faecalis* infection reduces lipid storage but increases the production of phospholipids that form the membranes of organelles, such as the endoplasmic reticulum, to support AMP synthesis^67^. These responses are context-and pathogen-dependent, reflecting the metabolic plasticity of the insect midgut. Immune pathway activation also appears to differentially regulates lipid storage across tissues and developmental stages^28,40,68^. Our results suggest that the midgut compensates by autonomously accumulating neutral lipids in response to enteric infection, potentially buffering systemic energy demand and supporting tissue-specific functions, such as lipophorin-mediated lipid trafficking and remodeling of tissue architecture, either through signaling or building cell structures.

While numerous lipid classes and species are implicated in immune modulation, our results highlight that a specific class with unique signaling properties (in a distinct tissue-type) may be crucial for broader host-pathogen responses. We found that 1,2-diacylglycerols are enriched in the midgut during enteric *P. e.* infection. 1,2-DAGs are important signaling messengers that play a role in regulating various physiological processes, including inflammation and muscle contraction^69^. During vertebrate immune responses, 1,2-DAGs are well-known as an important second messenger downstream of T cell activation through their T cell receptor (TCR)^70,71^. Furthermore, the absence of Diacylglycerol kinase ζ (DGKζ), an enzyme that converts DAG to phosphatidic acid (PA), leads to elevated DAG levels and prolonged DAG-mediated signaling, resulting in either enhanced immune activation or immune modulation depending on the immune cell type and infectious agent^72^. In addition, previous studies have uncovered that DAGs serve as a specific signal to initiate antibacterial autophagy through the activation of PKCδ during *S. Typhimurium* infection^73^. Notably, our work extends these findings by demonstrating that 1,2-DAG levels correlate with muscle contraction in the gastrointestinal tract of insects, a process that directly impacts pathogen expulsion. The specific regulation of 1,2-DAGs, as opposed to other lipid species like 1,3-DAGs or triacylglycerols (TAGs), underscores the cell-and tissue-type precision of lipid signaling in immune responses and suggests that subtle differences in the tuning of lipid metabolic cycles can profoundly influence physiological outcomes.

Host-pathogen interactions that shape gastrointestinal motility to either promote (or block) pathogen expulsion likely represent a conserved approach across taxa to control bacterial burden during enteric infection. Our results show that infection-induced signaling modalities (such PKC98e signaling, calcium signaling, and Tropomyosin1 [Tm1] function) within intestinal muscles are critical for defecation and bacterial pathogen clearance that impact infection outcomes, linking lipid metabolism and muscle remodeling during infection. In insects, the intestine acts as a primary barrier to oral infection, suggesting the maintaining midgut motility following enteric infection represents an ‘immune cell activation’ step that is crucial for host immunity. Indeed, work in Drosophila has highlighted that various excretion modules are critical for host-pathogen responses, including the removal toxic oxidized molecules that accumulate in various tissue-types in the host^74^.

Intestinal motility and pathogen clearance thus appear central to mucosal immunity, and our results suggest that distinct lipid metabolic adaptive responses are key to these physiological processes. Although our findings are built utilizing Drosophila models, DAG–PKC signaling pathways are evolutionarily conserved and regulate muscle contraction in vertebrates, including intestinal peristalsis^75^. This raises the possibility that similar lipid-regulated circuits operate in vertebrate intestinal physiology and/or immunity. In particular, our data support the concept that epithelial derived signaling lipids can influence tissue function non-autonomously, offering a blueprint for inter-cellular metabolic coordination of infection responses. Additionally, our study reveals that dietary fatty acid supplementation can significantly impact muscle function and enhance pathogen clearance, providing an avenue for dietary interventions in pathologies where gastrointestinal barrier function is compromised. While Drosophila offers a powerful genetic model of integrative physiology, further work is needed to validate these findings in vertebrate systems. Additionally, understanding how specific DAG species are generated and sensed across cells and tissues remains an open question. However, this study does broaden the conceptual landscape of lipid-mediated immunity and host-pathogen responses, as well as provides a foundation for exploring lipid-based interventions for enteric infection.

## Materials and Methods

### Drosophila husbandry and strains

The following strains were obtained from the Bloomington Drosophila Stock Center: *w1118* (#3605), *OreR* (#5), MexGal4 (#91368), ProsGal4 (#84276), NPFGal4 (#25682), HandGal4 (#66795), HowGal4 (#1767), Mhc-GFP (#38463), UAS-*Luciferase*^RNAi^ (#31603), UAS-*Bmm* (#76600), UAS-*Bmm*^RNAi^ (#25926), UAS-*Acc* (#63225), UAS-*Srebp*^RNAi^ (#34073), UAS-*Srebp* (#8236), UAS-*Lipin*^RNAi^ (#77170), UAS-*Mdy*^RNAi^ (#20167), UAS-*Lpr2*^RNAi^ (#31150), UAS-*ApoLpp* (#97113), UAS-*TrpA1*^RNAi^ (#31384), UAS-*DH31*^RNAi^ (#41957), UAS-*Itpr*^RNAi^ (#51795), UAS-*PKC98e*^RNAi^ (#35275), UAS-*PKC53e*^RNAi^(#55864), UAS-*inaE*^RNAi^ (#64885), UAS-*norpA*^RNAi^ (#31113), UAS-*PKCdelta*^RNAi^ (#28355), UAS-*aPKC*^RNAi^ (#25946), UAS-GCaMP6m (#42750), UAS-*Tm1*^RNAi^ (#43542). NP1Gal4 was kindly provided by D. Ferrandon.

All flies were reared on a standard yeast and cornmeal-based diet at 25 °C and 65% humidity on a 12 hr light and dark cycle, unless otherwise indicated. The standard lab diet (cornmeal-based) was made with the following protocol: 14g Agar, 165.4g Malt Extract, 41.4g Dry yeast, 78.2g Cornmeal, 4.7ml propionic acid, 3g Methyl 4-Hydroxybenzoate and 1.5L water. To standardize metabolic results, fifty virgins were crossed to 10 males and kept in bottles for 2-3 days to lay enough eggs. Progeny of crosses were collected for 3-4 days after initial eclosion. Collected progeny (males and females) were then transferred to new bottles and allowed them to mate for 3 days (representing unique populations fed *ad libitum*). All these flies were reared on a standard lab diet at 25 °C and 65% humidity on a 12 hr light and dark cycle, unless otherwise indicated. Next, post-mated females (20 flies per vial/cohort) were sorted into individual vials for natural oral infection.

Post-mated female flies were used for all experiments, because of sex-specific differences in *P.e.* infection sensitivity (males are significantly more resistant compared to females). All experimental genotypes were assayed for developmental defects (developmental timing, growth, fly size, and organ size), and no gross anatomical deficiencies were noted in any genotype represented in the results.

### Oral infection assays

*Pseudomonas entomophila* (*P.e.*) was used for natural (oral) infections^76^. Briefly, for oral infection, flies (app. 7 days of age, except for flies fed oleic acid supplemented food prior to infection) were placed in a fly vial with food/bacteria solution and maintained at 25 °C. The food solution was obtained by mixing a pellet of an overnight culture of bacteria (OD_600_= 50) with a solution of 5% sucrose (50/50) and added to a filter disk that completely covered the surface of standard fly medium. Mock treated flies were reared and cultured identically, only fed a 5% sucrose solution. Midguts of infected flies were dissected 20 h after oral contact with infected food. At least 3 vials (cohorts of 15 flies per vial) were used for each genotype for subsequent analysis. *P.e.* cultures are routinely genotyped for accuracy.

### Survival analysis

Flies were infected with *P.e* at OD_600_=50 as described above and kept at 25 °C for 20 hours. After overnight feeding (flies were always orally infected at 3:00-4:30pm), infected flies were transferred from vials with *P.e.* to vials containing standard lab food. Flies were transferred every day to a fresh vial for the first two days, and every two days after, and dead flies were counted every day until 8 days.

### RNA-Seq analysis

Intact fly (genotype: NP1Gal4>UAS-Lucif.^RNAi^) midguts (8) were dissected in PBS. Total RNA was extracted using Trizol reagent and used as template to generate sample libraries for RNA sequencing (Illumina Stranded mRNA Prep Kit). Sample libraries were sequenced using the Illumina MiSeq system. Sequence cluster identification, quality pre-filtering, base calling and uncertainty assessment were done in real time using Illumina’s HCS and RTA software with default parameter settings. Between 8 and 10 million (2X150) base pair reads were generated per library and mapped to the Drosophila genome (Release 6). Expression was recorded as TPM (transcripts per kbp per million reads) followed by Log2 transformation. Gene Ontology clustering analysis was performed using FlyMine. Currently, full completed analysis or RAW datasets will be available to reviewers upon request.

### Analysis of gene expression

Total RNA from intact fly intestines was extracted using Trizol and complementary DNA synthesized using Superscript III (Invitrogen). Real-time PCR was performed using SYBR Green, the Thermo Fisher QuantStudio 3 Real Time PCR system, and the primers pairs described in the extended experimental procedures (Table S1). Results are average ± standard error of at least three independent samples, and quantification of gene expression levels calculated using the △Ct method and normalized to actin5C expression levels (plotted as relative expression).

### Oleic acid feeding assay

For oleic acid (OA) supplementation diet, oleic acid (Sigma, # 364525) was incorporated into the standard lab food, resulting in 5% (w/v) concentration within food. Female flies were fed OA diet for 5 days, then conducting *P.e.* infection or dissection as needed.

### Food intake and feeding measurements

Twenty flies were transferred to vials filled with identical medium containing 0.5% brilliant blue (FD&C blue dye #1). Feeding was interrupted and 5 flies each were transferred to 200 ml 1× PBS containing 0.1% Triton X-100 (PBST) and homogenized immediately for 1 min by using pellet pestle mixer. Then samples were centrifuged at 4000 rpm and remove the pellet. Blue dye consumption was quantified by measuring absorbance of the supernatant at 630 nm (A630) by using Epoch plate reader (BioTek).

### Lipidomic and diacylglycerol analysis

Thirty whole Drosophila intestines including the anterior, middle, and posterior midguts (Genotype: OreR or NP1Gal4>UAS-Lucif.^RNAi^) were homogenized in 300 µL MilliQ water using a bead mill (25 Hz for 2 min). Following homogenization, an additional 650 µL water was added and a 20 µL aliquot was reserved for protein concentration. An aliquot of the homogenized sample (750 µL) was transferred to a glass tube containing methanol (900 µL) and an internal standard cocktail. Lipid extraction was performed by the addition of methyl-tert-butyl ether (3 mL) according to Matyash et al.^77^. Samples were injected onto an HPLC system connected to a triple quadrupole mass spectrometer (Sciex 5500QTRAP, Framingham, MA) and normal phase chromatography was performed using a HILIC column (2.1×100 mm, CORTECS 2.7 µm, Waters) ^78^. The mobile phase system consisted of solvent A (isooctane) and solvent B (isopropanol:isooctane:acetonitrile:water, 52:34:12:2, v/v/v/v) with 10 mM ammonium acetate and a post-column ionization buffer of isopropanol:water (95:5, v/v) with 30 mM ammonium acetate. Concentration was determined using stable isotope dilution with standard curves for saturated and unsaturated DAGs and results were normalized to protein content.

### Immunostaining and microscopy

Immunostaining was performed as described before^79^. Fly midguts were dissected in 1xPBS and fixed for 30 minutes at room temperature in gut fixation solution (100 mM Glutamic Acid, 25 mM KCl, 20 mM MgSO4, 4 mM Na2HPO4, 1 mM MgCl2; pH adjusted to 7.5 with 10 N NaOH, 4% Formaldehyde). Subsequently, all incubations were in PBS, 0.5% BSA, and 0.1% Triton-X at 4 °C. All primary antibodies were applied overnight. The following primary antibodies were used: rabbit anti-pan-PKC (ABclonal, #A17922, 1:500), mouse anti-Myosin heavy chain (MHC) (DSHB, # 3E8-3D3, 1:50). Fluorescent secondary antibodies (Jackson Immunoresearch, 1:500) and 40,6-diamidino-2-phenylindole (DAPI, 1:500) used to stain DNA was incubated at room temperature for 2h and 30 minutes, respectively. For visceral muscle staining, Alexa Fluor 555 Phalloidin (Thermo Fisher Scientific, 1:500) was used together with DAPI at room temperature for 30 minutes. Confocal images were collected using a Nikon Eclipse Ti confocal system (utilizing a single focal plane) and processed using the Nikon software and Adobe Photoshop. For quantitative analysis, images were analyzed by using ImageJ/Fiji software (https://imagej.net/software/fiji/).

### Bodipy staining and microscopy

Fly guts were dissected in PBS, then fixed in 4% formaldehyde/PBS for 20 min at room temperature. After washing with PBS for 10 min (3 times), guts were incubated in 2 μM Bodipy493/503 (Invitrogen #D3922), CholEsteryl Bodipy (Invitrogen #C12680) solution for 1h at room temperature. Then washed with PBS for 10 min (3 times), and incubated in DAPI (1:500) for 30 min. Guts were washed with PBS for 10 min (3 times), then mounted with Mowiol media. Samples were imaged by using Nikon Eclipse Ti confocal system (utilizing a single focal plane).

### Nile Red staining

Fly guts were dissected in PBS and fixed with 4% paraformaldehyde for 20 min at room temperature, washed 3 times with PBS for 10 min each, and then incubated with fresh Nile Red solution (2µl of 0.004% Nile Red Solution in 500 µl PBS) for 2 h at room temperature, followed by rinsing with PBS and then staining with DAPI (1:500). Confocal images were collected using a Nikon Eclipse Ti confocal system (utilizing a single focal plane) and processed using the Nikon software and Adobe Photoshop.

### RFP *P.e.* strain generation and oral infection

The DsRED-expressing strain of Pseudomonas entomophila was generated by introducing plasmid pSW002-Pc-DsRed-Express2 (Addgene, #11251) via electroporation^80^. *P. entomophila* was first grown in LB medium at 30 °C with shaking until reaching mid-log phase (OD600 =0.5–0.6). Cells were then harvested by centrifugation at 4 °C and made electrocompetent through successive washes with sterile distilled water followed by 10% glycerol. Electroporation was performed using a Bio-Rad electroporator (Micropulser™, Bio-Rad Laboratories) with a 0.2 cm gap cuvette. Fifty microliters of competent cells were mixed with 1 µL of plasmid DNA (approximately 100 ng) and transferred to the cuvette. A single pulse was applied at 2.5 kV. Following the pulse, cells were immediately recovered in 1 mL of antibiotic-free LB medium and incubated at 30 °C for 1–2 hours with shaking. Transformants were selected by plating on LB agar supplemented with tetracycline (15 µg/mL). RFP-*P.e.* infections were performed as described above.

### Measuring bacterial load (CFUs) in dissected adult midguts

Infected flies, dissecting forceps and dissecting dish were surface sterilized with 70% ethanol and washed with sterile PBS. Midguts from flies were then dissected individually in 1xPBS and each single gut was homogenized using a sterile pestle in 1.5 ml Eppendorf tube containing 200 µl 1xPBS. The homogenate was diluted to 1:1000 with 1xPBS. 50 µl of 1000-fold dilutions of the homogenate were plated onto LB plates and incubated at 29 °C overnight. The number of colonies in each plate were counted (only counting separated, well defined single colonies). At least 10 individual guts were measured in each treatment. Similar to bacterial cultures, these colonies derived from midgut dissections were routinely genotyped for accuracy and reproducibility of experiments (using PCR and primers specific for *P. e.*). All plates from mock-treated plate controls were negative for bacterial colonies.

CFU/fly for every plate was calculated using the following formulas:

CFU/ml = ((total number of colonies on a plate) * (dilution factor (1000)) / (plated volume: 0.05ml)

CFU/fly = ((CFU/ml) * total volume Eppendorf tube: 0.2ml)) / (number of flies per condition: 1 gut)

### Measuring defecation in adult flies

Defecation was measured by counting defecation ‘spots’ left dried on the inner wall of standard lab rearing vials. For the convenience of measuring defecation frequencies, 10 infected flies were transferred from vials containing bacterial solution (infection) to vials containing standard lab food with 1% Brilliant Blue FCF (no bacteria). The dried blue spots left on the inner wall of vials were counted after 16 hours. Experiments were repeated with at least 10 independent biological replicates/samples with 10 flies per replicate.

### Quantification and statistical analysis

Samples sizes were predetermined by the requirement for statistical analysis, and at least 3 biological replicates were utilized for all experiments. For all quantifications, n represents the number of biological replicates, and error bar represents SEM. Statistical significance was determined using either the unpaired Student’s *t*-test or two-way ANOVA with Tukey post hoc test where multiple comparisons were necessary, in GraphPad Prism Software, and expressed as *P* values.

## Acknowledgments

This work was supported by the National Institute of Diabetes and Digestive and Kidney Diseases (grant R01DK133294 to J.K.). The authors wish to thank the Nutrition Obesity Research Center (NORC) Lipidomics Core Laboratory (P30DK048520) at the University of Colorado Anschutz Medical Campus for their contributions to this manuscript.

## Author contributions

X.L. designed and performed experiments, as well as wrote the manuscript. M.M. generated RFP-*P.e.* strain. J.K. designed experiments and wrote the manuscript.

