## Supplemental Files for "Diacylglycerol metabolism drives host-pathogen responses during enteric infection in *Drosophila*"

**Table S1: qRT-PCR Primers**

| Primer | Sequence (5’-3’) |
|---|---|
| actin5C-F | CTCGCCACTTGCGTTTACAGT |
| actin5C-R | TCCATATCGTCCCAGTTGGTC |
| Fasn1-F | AGCTAATAACGGCAGTCAACGG |
| Fasn1-R | CAGGTTTAGTTGTAGGGGCTAGA |
| Lipin-F | GTCCGGTACGAAGAAGTCCG |
| Lipin-R | TCTGAGATACGGCAACTGCT |
| Mdy-F | ACACAAAGTTCCCAGAGTTCA |
| Mdy-R | TGGTGATGGTCTCGATTGGA |
| Drs-F | AAGTACTTGTTCGCCCTCTTCGCT |
| Drs-R | TCCTTCGCACCAGCACTTCAGACT |
| Dro-F | TTCACCATCGTTTTCCTGCT |
| Dro-R | GGCAGCTTGAGTCAGGTGAT |
| CecA-F | CATTCTGGCCATCACCATTGGACA |
| CecA-R | ACATTGGCGGCTTGTTGAGCGATT |
| Def-F | GAGCCACATGCGACCTACTC |
| Def-R | CAGTAGCCGCCTTTGAACC |
| AttA-F | TCGTTTGGATCTGACCAAGGGCAT |
| AttA-R | TTCCGCTGGAACTCGAAACCATTG |

### Supplementary Figures

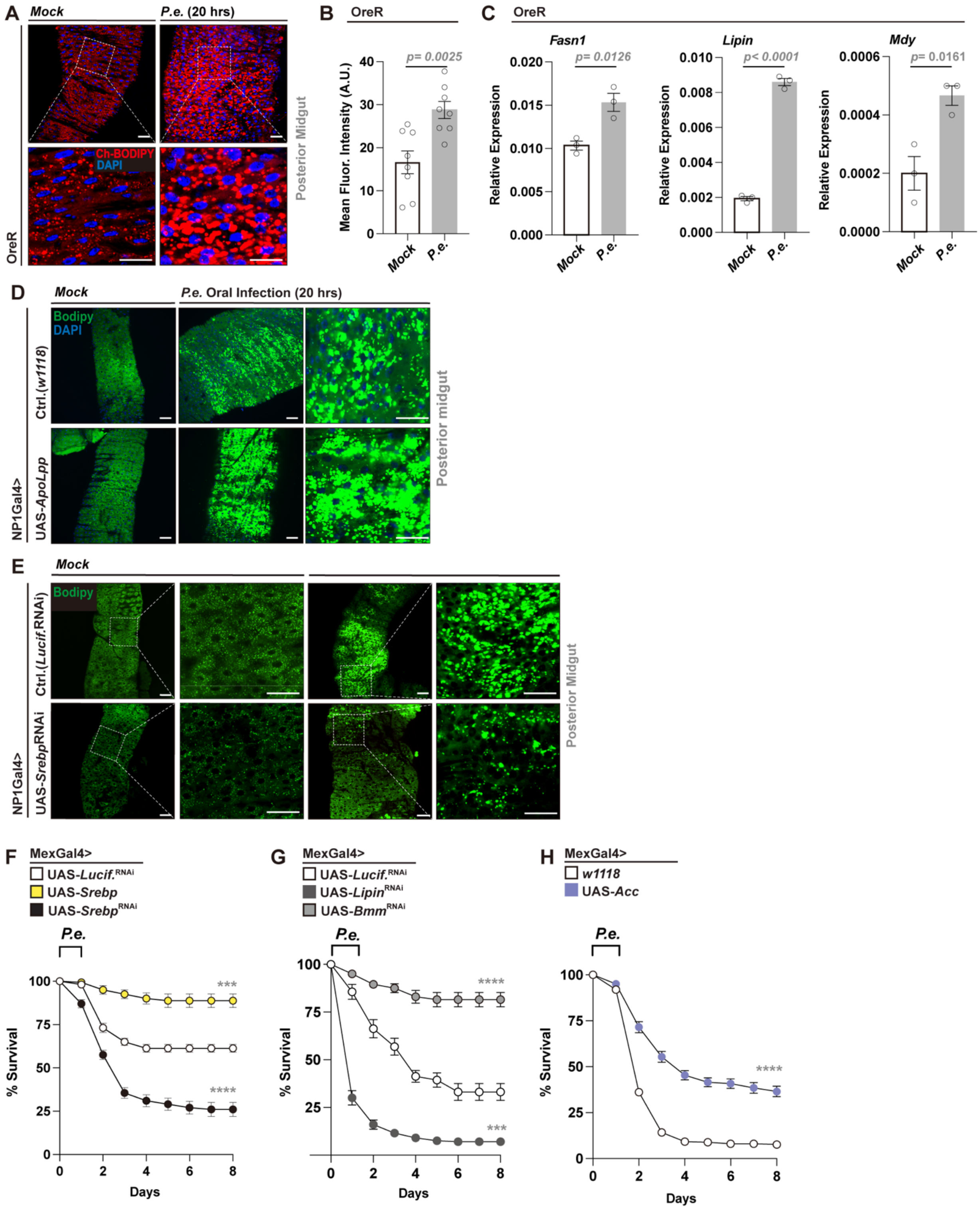


**Figure S1. *P.e.* infection alters lipid metabolism in the Drosophila midgut, and enterocyte lipid homeostasis influences host survival post *P.e.* infection.** (A, B)

CholEsteryl BODIPY (Ch-BODIPY, red) staining (A) and quantification (B) of cholesterol levels in the posterior midgut of OreR flies during mock or P.e. infection (n = 8 midguts). Nuclei (DAPI, blue). (C) qRT-PCR validation showing *P.e.* infection-induced upregulation of genes involved in lipid anabolism. (D) Enhanced lipid accumulation in NP1Gal4>UAS-*ApoLpp* posterior midgut following infection, revealed by BODIPY 493/503 staining (green). (E) Suppressed lipid accumulation in NP1Gal4>UAS-*SREBP*$^{RNAi}$ flies during infection, as indicated by BODIPY 493/503 staining (green). (F–H) Survival analysis of flies with EC-specific genetic manipulation of lipid metabolism using NP1Gal4 or MexGal4 drivers. Attenuation or overexpression of *Srebp*, *Bmm*, *Lipin*, or *Acc* significantly alters survival compared to NP1Gal4/MexGal4>UAS-*Lucif.*$^{RNAi}$ controls (n = 200 flies/genotype). Statistical comparisons were performed using two-sided t-tests (B, C) or two-way ANOVA (F-H). Bars represent mean ± SEM. ***p value < 0.001, ****p value < 0.0001. Scale bars, 10 μm.

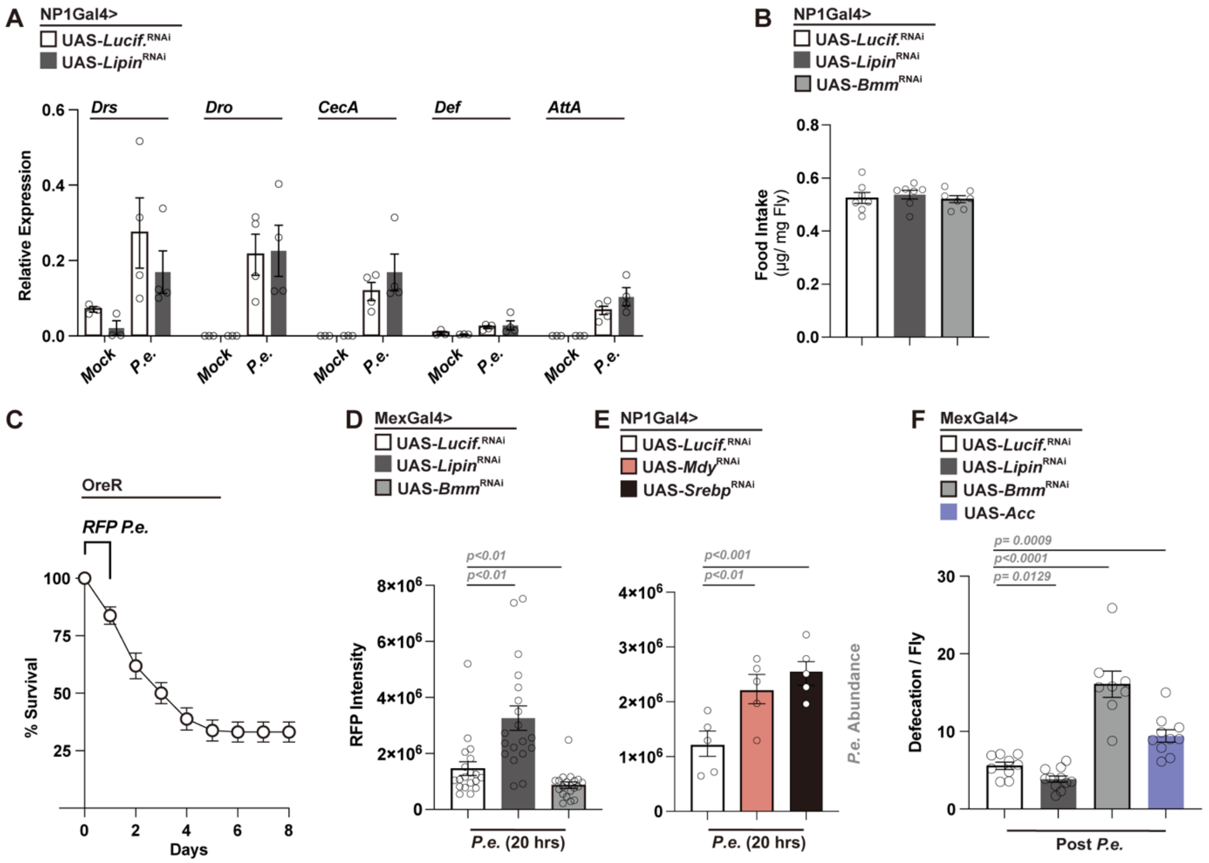


**Figure S2. Enterocyte lipid metabolism regulates host survival by modulating gut motility and bacterial clearance.** (A) Expression of antimicrobial peptide genes (*Drs, Dro, CecA, Def, Atta*) in midguts of control and *Lipin*$^{\text{RNAi}}$ flies shows no significant difference post *P.e.* infection. (B) Food intake remains unchanged in flies with EC-specific attenuation of *Lipin* or *Bmm*. (C) DsRed-expressing *P.e.* (RFP-*P.e.*) is pathogenic and influences host survival after infection. (D, E) EC-specific attenuation of lipid metabolic genes affects bacterial clearance during RFP-*P.e.* infection. (F) Defecation frequency is altered in flies with EC-specific modulation of lipid metabolism. Data represent mean ± SEM. Statistical comparisons were performed using two-sided *t*-tests (A, B, D-F).

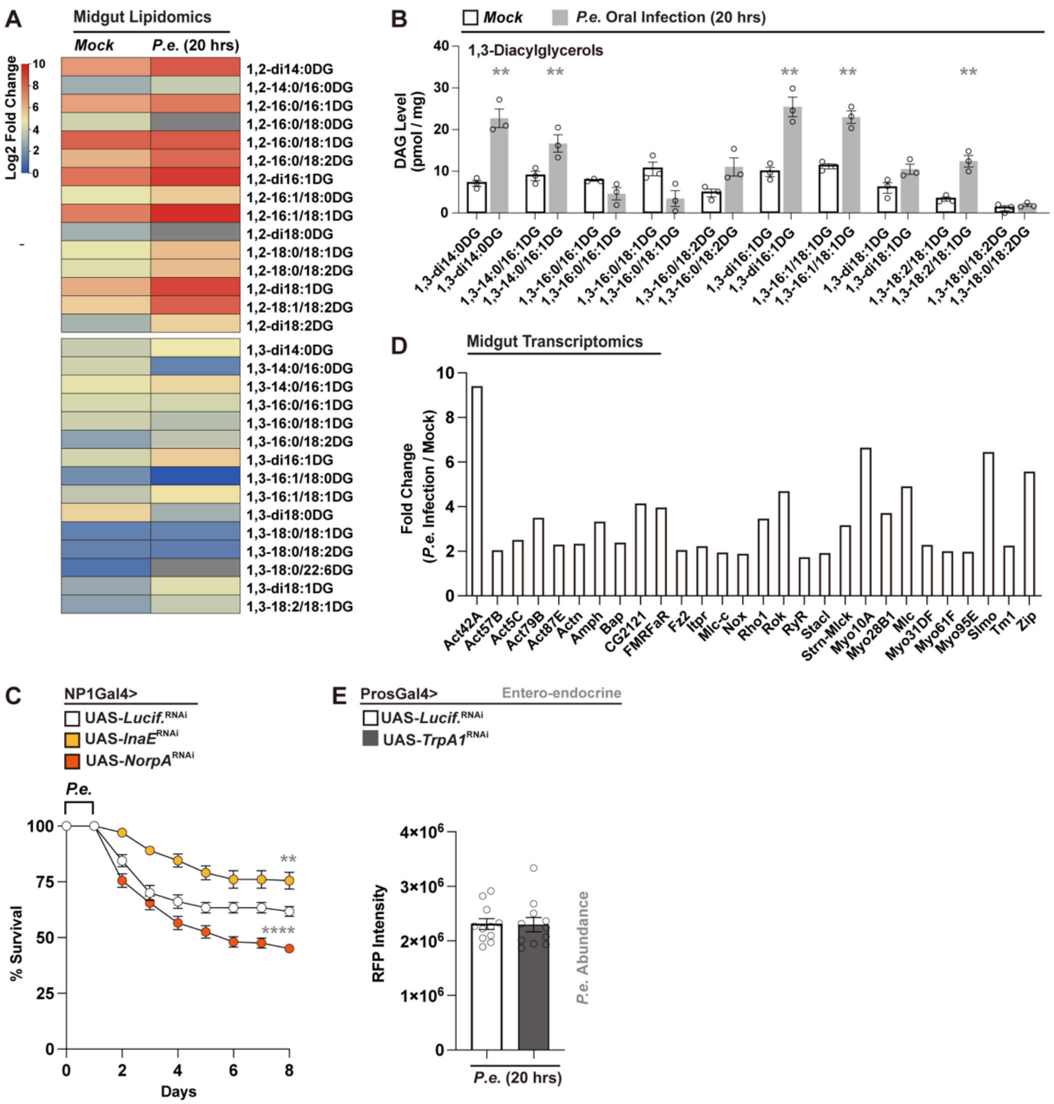


**Figure S3. *P.e.* infection induces diacylglycerol (DAG) accumulation and activates visceral muscle remodeling.** (A, B) Heatmap (A) and quantification (B) of midgut lipidomic profiles reveal specific elevation of DAGs post *P.e.* infection. Heatmap representation of lipid species in dissected midguts from control (Mock) and *P. e.*–infected flies (20 h post-infection). Genotype: NP1Gal4>UAS-Lucif.$^{RNAi}$. (C) EC-specific attenuation of *InaE* or *NorpA* alters survival following infection. (D) Gut RNA-seq analysis identifies infection-induced upregulation of muscle contractility-related genes. (E) EEC-specific attenuation of *TrpA1* does not affect bacterial clearance post *P.e.* infection. Statistical comparisons were performed using two-sided t-tests (B, E) or two-way ANOVA (C). Bars represent mean ± SEM. **p value < 0.01, ****p value < 0.0001.

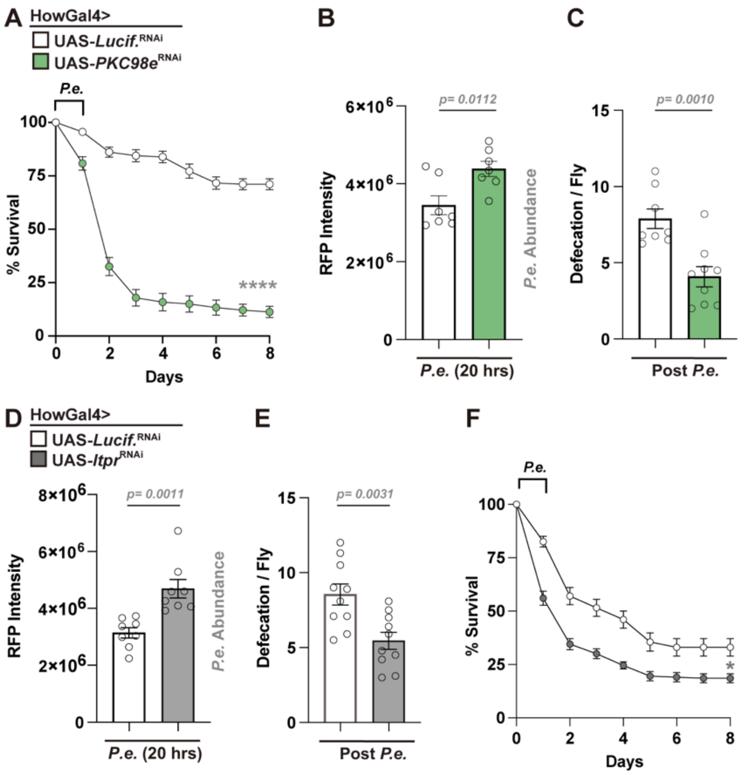


**Figure S4. Visceral muscle PKC–calcium signaling regulates muscle contraction, pathogen clearance, and host survival.** (A–C) Attenuation of PKC98e in visceral muscle (HowGal4>UAS-*PKC98e*$^{\mathrm{RNAi}}$) reduces survival (A), impairs bacterial clearance (B), and decreases defecation (C) post-infection. (D–F) Visceral muscle-specific attenuation of Itpr (HowGal4>UAS-*Itpr*$^{\mathrm{RNAi}}$) similarly compromises bacterial clearance (D), reduces defecation (E), and lowers survival (F). Statistical comparisons were performed using two-sided *t*-tests (B-E) or two-way ANOVA (A, F). Bars represent mean ± SEM. *p value < 0.05, ****p value < 0.0001.

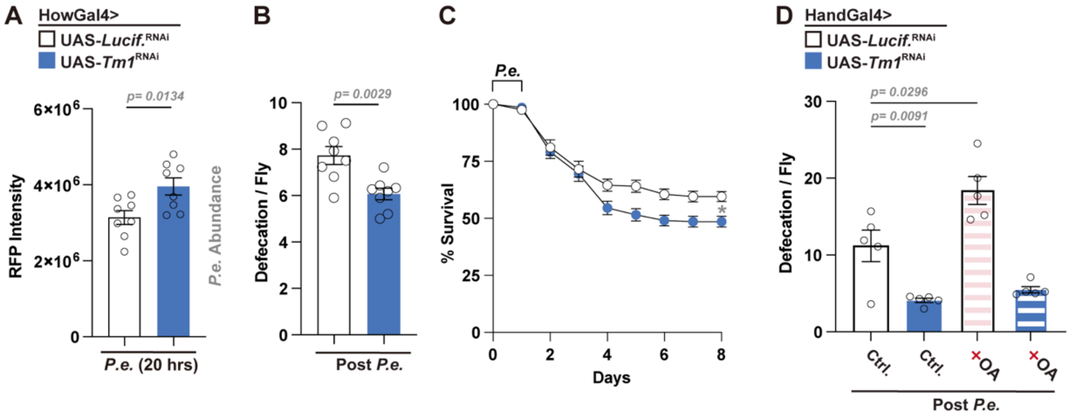


**Figure S5. Tropomyosin1 (Tm1) is required in visceral muscle for contraction, pathogen clearance, and host survival.** (A, B) Attenuation of Tm1 in visceral muscle (HowGal4>UAS-*Tm1*^RNAi^) increases midgut bacterial load (A) and reduces defecation frequency (B) post *P.e.* infection. (C) Survival curves showing decreased survival in *Tm1*-deficient flies compared to controls. (D) Dietary OA supplementation prior to infection partially restores defecation in HandGal4>UAS-*Tm1*^RNAi^ flies. Statistical significance determined using two-tailed t-tests (A, B, D) or two-way ANOVA (C). Bars represent mean ± SEM. *p value < 0.05.
